# Sensorimotor dynamics in the superior colliculus of the echolocating bat

**DOI:** 10.1101/2025.08.26.672231

**Authors:** Gowri Somasekhar, Ninad B. Kothari, Cindy F. Moss, Melville Wohlgemuth

## Abstract

To successfully execute natural tasks, animals must continuously monitor and integrate dynamic sensory information to guide behavior. While this process – known as active sensing – is fundamental to goal-directed behavior, the mechanisms by which sensory information supports motor planning, particularly across different behavioral contexts, remains poorly understood. We investigated sensory and motor activity in the superior colliculus (SC) of echolocating bats, a powerful model system to study active sensing behaviors due to their reliance on self-generated sonar signals. We recorded multichannel SC activity while bats performed spatial navigation and target tracking behaviors. We hypothesized that changes in sensory signals (i.e., returning echoes) drive quantifiable adjustments in SC activity related to sonar call production, and that adaptive changes in SC sensorimotor processing are preserved across behavioral contexts. We examined pooled SC activity using a dimensionality reduction technique to isolate the largest changes in firing rates across multichannel SC recordings. We then examined the variation in SC activity from the time of echo arrival (sensory input) to call production (motor output). Our data show that increases in call rate are associated with shorter trajectory lengths in neural state space, reflecting reduced variability and increased efficiency in population activity. Strikingly, when bats cluster calls into sonar sound groups (SSGs, which are rapid bursts of calls to increase sensory sampling), SC activity is distinct from times when bats produce isolated calls. Notably, we find that successive SSGs lead to progressively shorter trajectory lengths, suggesting a refinement of sensorimotor processing across successive SSGs. Together, our findings demonstrate that sensorimotor population dynamics in the SC follow common principles across distinct behavioral tasks. These results support a fundamental role for the SC in adaptive sensing and highlight its contribution to flexible sensorimotor tranformations in natural behaviors.

## Introduction

In the natural world, animals must contend with complex and dynamic stimuli, which has shaped the evolution of sensorimotor systems that filter and select behaviorally relevant stimuli. For example, primates move their eyes to scan and foveate objects in the environment (Nelson and MacIver, 2006; Schroeder et al., 2010). Echolocating bats, our model animal, modulate their sonar calls to refine spatial information carried by echo returns and to focus on objects (Griffin, 1958; Griffin et al., 1960). These active sensing behaviors operate as closed-loop systems, whereby sensory feedback informs subsequent motor commands (Yang, Wolpert and Lengyel, 2016). How sensing influences upcoming motor commands is still a topic of active research and the focus of the current study.

The echolocating bat is a powerful model system to probe the temporal dynamics of sensorimotor integration, as echo reception and call production are temporally discrete and measurable events, providing direct and accurate metrics of sensing and action on a millisecond time scale. Bats produce sonar calls and listen to returning echoes, rapidly adapting sonar call features, including adaptive changes to the timing of calls, in response to information conveyed in echoes (Griffin, 1958; Luo et al., 2017; Moss et al., 2006; Moss et al., 2011; Petrites et al., 2009). As a common active sensing strategy while tracking targets both in flight and from stationary positions, bats increase sonar call rate, reducing the interval between calls, as the distance of the target decreases (Simmons et al., 1979; Kalko and Schnitzler, 1989; Moss and Surlykke, 2001, Aytekin et al, 2010, Kothari et al, 2014). In addition to modulating call rate, bats also dynamically control the timing of their calls producing clusters of sonar calls, called sonar sound groups. Such temporal grouping of sonar calls is thought to be essential for extracting task relevant information from the environment (Moss and Surlykke, 2001, Moss and Surlykke, 2010) and has also been found to be a common active sensing strategy displayed by bats across behavioral contexts—such as in the presence of clutter (Petrites et al, 2009, Moss et al, 2006), while tracking eratically moving prey in-flight (Moss and Surlykke, 2010) as well as from a stationary position (Aytekin et al, 2010, Kothari et al, 2018), or with increasing task difficulty (Sandig et al 2014, Petrites et al, 2009). While both temporal rate adjustment and production of sonar sound groups are shared active sensing strategies across behavioral contexts, it is unknown whether the underlying neural correlates are also common across behavioral contexts or different.

In the current study, we investigated neural population activity in the time window between sensory processing (echo arrival) and motor action (call production). Our goal was to quantify changes in population-level brain activity in this sensorimotor window under different task demands (navigation versus pursuit/tracking) and changing sensory conditions. We further asked whether active sensing strategies which are shared across different behaviors, such as sonar sound group production and call rate modulation, are accompanied by shared neural signatures in the brain, potentially revealing generalizable principles of sensorimotor integration across behaviors.

Here, we focused on the midbrain superior colliculus (SC), a major hub of sensorimotor processing for spatial orientation and target selection (Gandhi and Katani, 2011, Basso et al, 2021, Allen et al, 2021). The SC has been shown to serve an integral role in spatial attention networks and drives species-specific orienting behaviors (Knudsen, 2007, Krauzlis et al, 2013, Allen et al, 2021). The SC receives task- and context-specific inputs from the cortex, which are believed to influence attention for target selection and decision making (Knudsen, 2007, Krauzlis et al, 2013). The SC also receives inputs from early-stage sensory processing areas that can influence motor planning on a more rapid timescale (Krauzlis et al, 2013). Studying combined SC neural activity therefore affords an opportunity to examine how sensory information arriving from multiple circuits influences outgoing motor signals. The sensing behaviors of the echolocating bat are well suited for this study because sensory (echo) and motor (call) events are temporally precise and segregated in time; and sensory and motor signals for echoes and calls, respectively, were found in the bat SC in past work (Valentine and Moss, 1997, Wohlgemuth and Moss, 2016, Kothari et al, 2018). Moreover, the bat makes measurable and predictable changes in sonar signal design based upon arriving echo information, providing a quantifiable metric (e.g., changes in call rate, discussed below) to analyze sensorimotor processing with respect to adaptive behavioral control.

In this study, we analyze multichannel neural activity across the SC of echolocating bats performing navigation and target-tracking tasks. To probe population dynamics, we leveraged a dimensionality reduction-based approach developed for motor and premotor cortical analysis during sequenced behaviors (Bernardin et al., 2012; Churchland et al., 2006, 2012; Russo et al., 2020; Shenoy et al., 2013; Zimnik and Churchland, 2021). These frameworks capture how activity across neurons evolves across different task periods, such as from motor preparation to execution (Shenoy et al., 2013; Zimnik and Churchland, 2021). In this study, we analyzed the bat’s context-dependent adjustments in call rate to test the hypothesis that changes in sensory input (i.e.,returning echoes) elicit measurable modulations in SC premotor activity for vocal output. Specifically, we predict that SC population activity occupies distinct neural subspaces corresponding to key epochs: echo arrival, sensorimotor integration, and motor planning for the subsequent call. Additionally, we predict that SC activity patterns diverge systematically when the bat modulates the temporal structure and rate of sonar calls in response to varying environmental demands.

## Methods

### Animal subjects and training

Data from five adult big brown bats (*Eptesicus fuscus*) are included in this study. The bats were collected in the State of Maryland under a permit issued by the Department of Natural Resources and were housed in animal vivariums at the University of Maryland and Johns Hopkins University. All protocols and procedures were approved by the Institutional Animal Care and Use Committees (IACUC) at both institutions, where this research was conducted.

We trained two bats to perform a spatial navigation task, referred to as the free-flight task, in which they freely flew in a large experimental flight chamber (6x6x2.5 meters), to land on a platform or avoid stationary obstacles (see Fig 1a and Kothari et al., 2018 for complete details). In both versions of this task, the bats identified object locations mid-flight and adjusted their sonar and flight behaviors accordingly. In a second task, three other bats were studied in a sonar target tracking paradigm, termed the *platform-tracking task*, where they tracked and intercepted a moving insect from a stationary position on a platform (see Fig 1b, and Wohlgemuth et al., 2018 for complete details). In this paradigm, a tethered insect was propelled towards the bat using a computer-controlled stepper-motor, and the movement of the target was under experimental control. For both tasks, brain recordings were collected using a 16-channel silicon probe (Neuronexus) arranged in a 4x4 grid, with 125-micrometer spacing between sites. The silicon probe was mounted on a moveable microdrive, which permitted sampling of neural activity along the dorsal-ventral axis of the SC. For the free-flight task, the recordings were made with a wireless telemetry system (TBSI, Triangle Biosystems International) that was connected to a neural acquisition system (Plexon). The neural recordings in the platform-tracking task were made using a tethered system, also from Plexon. All neural recordings were digitized at a sampling rate of 40 kHz.

**Figure 1.**
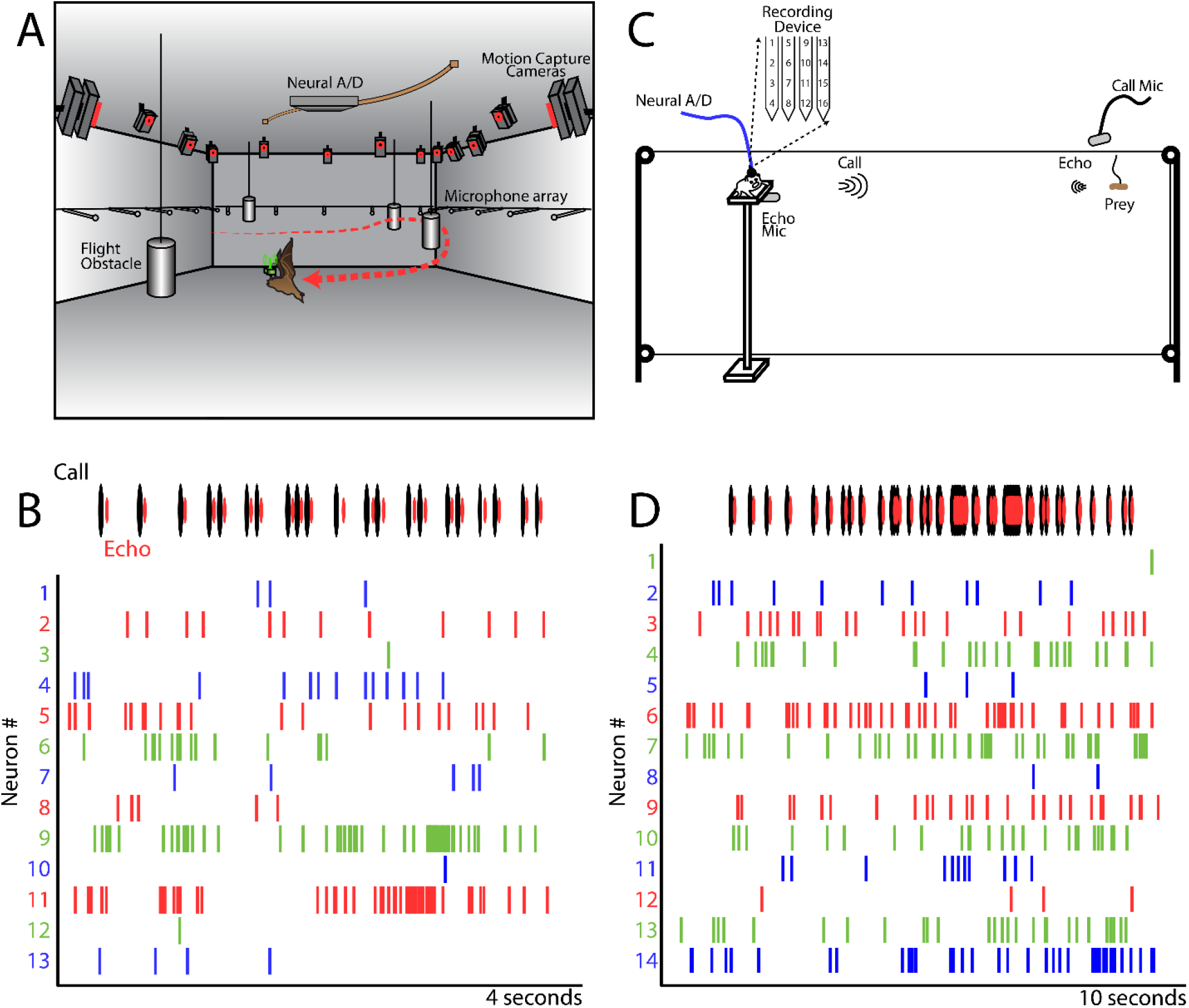
A.Illustration of the free-flight task. The bat is tasked to fly around in a cluttered environment and avoid crashing into obstacles. While the bat performs these behaviors, microphones record the bat’s calls, motion capture cameras record its position relative to objects, and a wireless recording system collects brain activity. B. Top, call (black) and echo (red) times for one trial from a recording session where the bat navigated around the flight chamber and avoided obstacles. Bottom, example raster plot of the activity of 13 neurons recorded during the same flight as the microphone recordings. Neuron identity is indicated along the vertical axis, with tick marks along the horizontal axis indicating the spike times for each neuron on this trial. C. Illustration of the platform-tracking task. The bat is trained to track a moving prey item while it remains stationary on a small platform. The movement of the target is controlled by stepper motors, sonar call and echo recordings were made with microphones, and neural recordings collected with a tethered physiology system. D. Top, call (black) and echo (red) times for one trial of the bat tracking a moving target. Bottom, raster plot of 14 different neurons’ activity during the same trial as the microphone recordings.

### Data collection

Behavior was quantified with high-speed video recording and audio acquisition systems. For the free-flight task, we recorded the bat’s sonar calls on a 32-channel wideband microphone array. The microphones were evenly spaced (∼1 meter apart) in a horizontal plane, 1.5 meters from the floor of the room. The microphone array recorded the bat’s calls as it flew in multiple directions around the experimental room. The microphone array was also used to reconstruct the sonar beam direction and width (Ghose and Moss, 2003; Jakobsen et al., 2015; Kothari et al., 2018; Surlykke et al., 2009) and to compute the time of echo arrival at the bat’s ears. Bat echolocation calls were filtered (10-100 kHz) and digitized with a 32-channel A/D board (National Instruments) using a sampling rate of 250 kHz per channel. The bat’s 3D flight trajectory, head aim, and location of obstacles were tracked with a high-speed 16-camera motion capture system (Vicon) at a sampling rate of 300 fps. On each experimental day, the animal was released from multiple locations in the room to cover the entire flight space of the chamber. Synchronization across recording systems was done with a common end trigger manually initiated each time the bat landed after flight.

Audio and neural data were collected similarly in the platform-tracking task (Fig 1c). The bat’s calls were recorded with an ultrasonic microphone placed in front of the animal at 3 meters, and returning echoes were collected with a microphone placed below the platform where the animal was seated. Audio data were filtered (20-100 kHz) and digitized at a sampling rate of 250 kHz per channel. Motion capture data were not taken in the target tracking task, because the bat remained in a stationary position, and the target’s movement was controlled by a computer and therefore at known distances from the bat during the experiment. A dog clicker training device prepared the bat for each trial, and then the stepper-motor movements were initiated by a keystroke on a computer (see Wohlgemuth et al., 2018 for complete details). The target then moved towards the bat following a variety of pre-programmed trajectories while the bat tracked its position using sonar. Synchrony across the neural, audio and target trajectory control systems was achieved with a common start trigger generated by the computer controlling the stepper motors and then sent to the other hardware systems.

### Data analysis

The neural and behavioral data were processed *post hoc* using motion analysis software (Vicon Nexus) and custom code written in Matlab. In the free-flight task, motion capture data was analyzed to extract the head aim of the bat, the 3D position of the bat, and the 3D positions of the flight obstacles. Three reflective markers affixed to the top of the telemetry board (on the animal’s head) were used to construct the head-aim direction vector, and the 3D position of the bat (Kothari et al, 2018). Similar markers were used to keep track of the locations of objects in the room. Using this data, we calculated the egocentric location of objects within the bat’s *acoustic field of view* (Jakobsen et al., 2013), and then determined the timing of echoes returning from objects to the bat’s ears for each emitted call.

Audio recordings were first processed to extract the onset and offset times of each sonar call. In the free-flight experiment setup, multiple microphones were mounted on each wall, and we selected the microphone for analysis based on the bat’s flight and head aim (i.e., we chose the microphone the bat flew towards for maximum SNR recordings). In the platform target-tracking task, the bat faced the moving prey, and call data was taken from a microphone facing the bat 3 meters from the platform. The timing of all calls was corrected by the distance between the bat and the microphone.

After determining the call onset and echo arrival times, we analyzed call intervals (i.e., the interval from the onset of one sonar call to the next) and whether the bat was producing sonar sound groups (SSGs). SSGs have been described in prior reports (Hulgard and Ratcliffe, 2016; Kothari et al., 2014; Moss and Surlykke, 2001), but in brief, an SSG is defined as a cluster of two or more sonar calls at comparatively short and regular intervals, flanked by single calls produced at longer intervals. When comparing neural activity for SSG and non-SSG calls, we restricted the range of call intervals to 40-100 milliseconds to control for any possible call interval effects on neural activity patterns. Within a trial, data (behavioral and neural) were used from periods when the bat was ‘engaged’ in the task. For the recordings taken in the free-flight obstacle avoidance task, this meant only times when the bat was in flight, and not the pre- and post-flight times. For the platform tracking task, we used data collected during the movement of the target (i.e., not before and after movement), and only included trials where there was a decrease in call interval (i.e., increase in the rate of calls) as the target approached the bat.

The neural recordings were first processed using a wavelet-based clustering algorithm to sort the units recorded across the 16-channel probe in each experimental session (Kothari et al., 2018; Quiroga et al., 2004; Wohlgemuth et al., 2017). Once the spike times of individual units were determined, we then analyzed neural data with respect to audio data. We wanted to examine neural activity in the time window beginning at sensory processing and ending at the beginning of the next motor command – for the bat, this window begins at the arrival time of an echo and extends to the onset of the next call. For instance, if there were 100 sonar calls and echoes in one particular trial, the first window would start at Echo-1 arrival and end at Call-2 onset, then the next window would start at Echo-2 arrival and end at Call-3 onset, then Echo-3 arrival to Call-4 onset, and so on until the end of the trial, resulting in 99 echo-to-call windows. This procedure was performed for all sites where at least three neurons were recorded simultaneously.

Once the echo-to-call aligned windows of neural activity were extracted, several different analyses were performed to construct the matrices for dimensionality reduction. First, we warped the length of each echo-to-call neural window to the mean window length for that experimental session (see Fig 3; Leonardo, 2004; Sober et al., 2008). This is necessary because the length of time between echo arrival to the onset of the next sonar call varies with the bat’s call rate and echo arrival times. As such, it is not possible to combine across ‘unwarped’ echo-to-call windows because they are all different lengths of time. The warping procedure is similar to those used in birdsong studies when the lengths of syllables vary from rendition to rendition and spike times are warped to compare average activity patterns over renditions (Leonardo, 2004; Sober et al., 2008). For the current data, the windows were warped by first finding the mean length of time from echo arrival to the next call onset across an experimental session. Then, spike times in the unwarped neural windows were calculated (i.e., the actual spike times), and then converted into percentages of time through the window. These percentages were then used to warp the spike times into a window size that matched the mean length of all echo-to-call windows. Once these warped spike matrices were constructed, we then calculated instantaneous spike rates from the warped spike times. These spike rates were then z-scored, and a principal component analysis (PCA) was performed on the z-scored spike rates.

Once the PCA dimensionality reduction was done, we examined the PCA scores with respect to differences in the bat’s behavior. The first three principal components were used for all analyses. PCA scores were then averaged within defined behavioral conditions. For instance, when comparing the coordinated neural activity for different ranges of call intervals for the platform tracking task, we averaged the PCA scores across all echo-to-call windows in 50 millisecond bins from 0-200 milliseconds (i.e., 0-50, 51-100, 101-150, and 151-200 msec). For the free-flight task there was a smaller range of call intervals, so 3 bins between 0-90 milliseconds were used. We then compared the lengths of the neural trajectories across each call interval bin by summing the distances between successive points in the averaged 3-dimensional vector. More specifically, for each 3-dimensional vector we calculated the distance from point 1 to point 2, point 2 to point 3, and so on for the entire length of the vector. We then summed those distances to get a total 3-dimensional vector length. This was done across all sites in all bats for both the free-flight and platform-tracking tasks.

As a control analysis, we performed a randomization procedure to select random start times for neural windows of interest within each day’s recording session. For the random data set, we chose the same number of randomly selected windows as in the original data set. As an example, if there were 500 echo-to-call windows with intervals less than 50 msec, then the randomly selected data set would also have 500 randomly selected windows of 50 msec duration (and an equal number of windows across all call interval bins). The random start times were chosen by taking the time series of the original neural data and randomly selecting a number from that time series as the beginning of the random window. The goal of the random selection was to have a comparable selection of data to the original data set, but where the start time of each window is not associated with echo arrival times. Once these randomly selected windows of neural activity were extracted from the data, the same time-warping and PCA-based measures were used as those applied to the original data set for each recording session. Statistical analysis to compare the differences in trajectory lengths between call-interval bins and between the original and random data sets were performed using a 2-way, repeated measures ANOVA.

A similar process was performed to compare differences in echo-to-call windows for SSG and non-SSG production. SSGs are windows of time when the bat increases sensory sampling and homing in on a target or goal (Falk et al., 2014; Kothari et al., 2014; Petrites et al., 2009; Sändig et al., 2014). We wanted to examine how neural activity is influenced by this period of increased sensory acquisition. We averaged PCA scores in echo-to-call windows for SSGs and non-SSGs. We then measured the distances between the SSG and non-SSG trajectories using a 3-dimensional Euclidean distance measurement in a point-by-point manner. For example, if the SSG and non-SSG trajectories were 100 points in length, we calculate the distance from point-1 SSG to point-1 non-SSG and so on for all 100 points along the two vectors. This would then generate a 100-point vector of distances between the SSG and non-SSG trajectories for this example. For the analyses of successive SSG production (Fig 9), we examined the length of trajectories by calculating the Euclidean distances between successive points in each 3D vector (e.g., similar to the trajectory lengths calculated for different echo-to-call window lengths, Fig 5 and 6). A permutation test was used to test for significant differences across SSG and non-SSG neural trajectories.

## Results

We investigated neural population dynamics in the superior colliculus (SC) from the moment of sensory acquisition (echo arrival) to motor initiation (sonar call production). We also compared sensorimotor processing across distinct behaviors to identify common neural dynamics. To do this, we studied five bats across two tasks. We trained two bats on a free-flight task, where they flew through a large experimental chamber and navigated stationary obstacles (see Kothari et al.,2018; Fig. 1a). We trained three other bats on a platform-tracking task, where they tracked a moving target while remaining stationary on a platform (see Wohlgemuth et al., 2018; Fig. 1c). While the bats performed these tasks, we recorded their echolocation calls, tracked the 3D positions of the bat and objects, and monitored neural activity in the SC (see Fig. 1b and 1d for examples of multi-channel recordings). All behavioral and neural data is time-synchronized to align brain activity to changes in behavior after data collection.

In the two bats that performed the free-flight task, we recorded from 181 neurons; and in the three bats that performed the platform-tracking task, we recorded from 463 neurons. The 16-channel recording device allowed us to sample more than one neuron simultaneously during each recording session. In the free-flight task, 9 different recording sessions were performed with multiple neurons (mean 10.7 neurons/session); and in the platform-tracking task, 45 different sessions had multiple neurons recorded simultaneously (mean 10.3 neurons/session).

This study investigated the relationship between sensory coding and signals for adaptive motor control. Our recording device spanned multiple layers of the SC, and our recordings therefore contain a mix of sensory and/or motor signals because neurons of each functional class can be found throughout the bat SC (Wohlgemuth et al., 2018; Wohlgemuth and Moss, 2016). We characterized changes in population coding of sensory-to-motor events in the SC using a dimensionality reduction approach to identify the largest changes in spiking activity across simultaneously recorded neurons (Churchland et al., 2012). The data from all recorded neurons from each recording session were collated and separated into time segments beginning with echo arrival and ending with sonar call onset (see Fig 2a). These windows therefore define the time period over which sensory signals change into motor commands in the SC. We then characterized changes in coordinated activity patterns across neurons within these echo-to-call windows.

**Figure 2.**
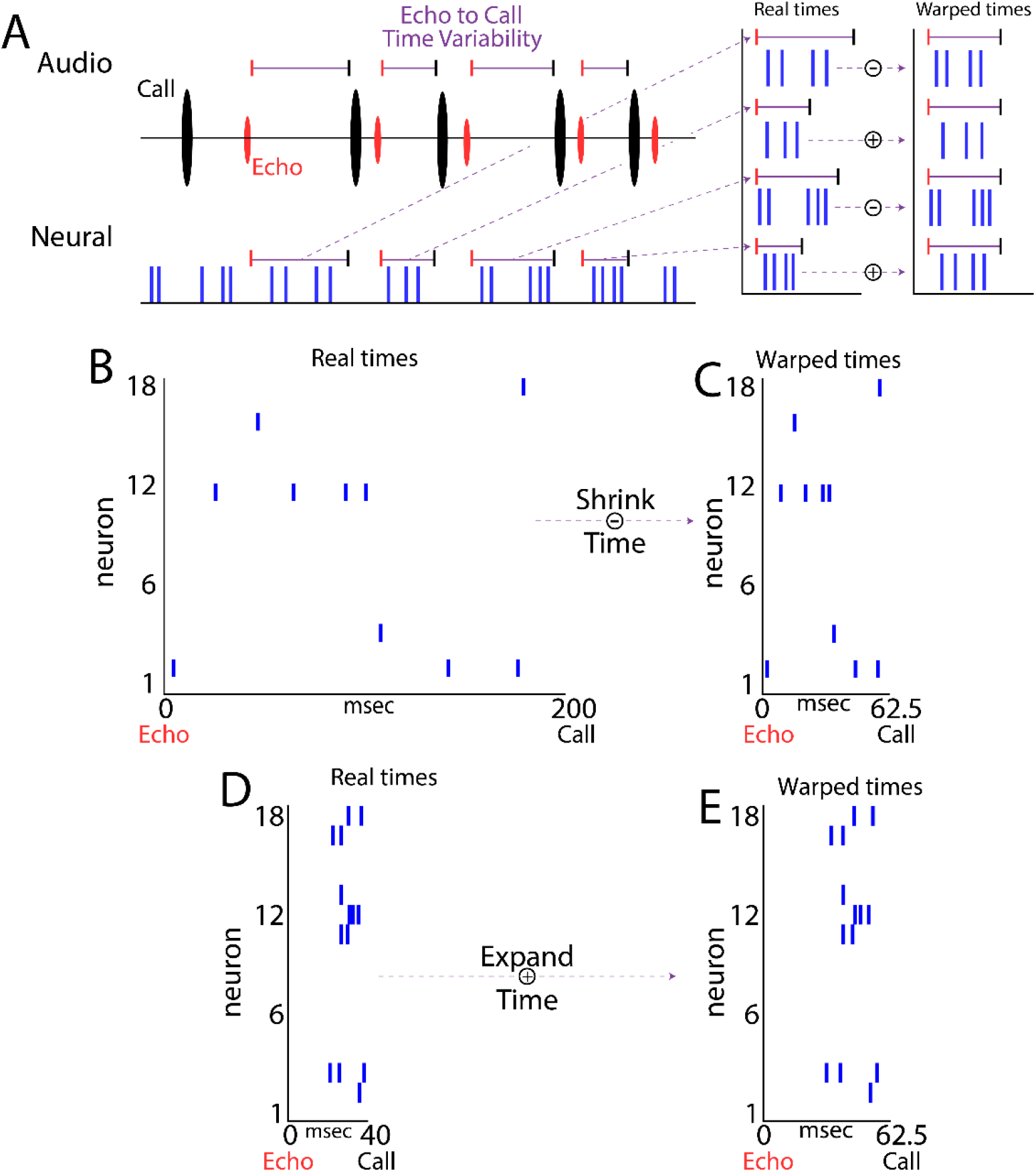
A.Schematic of the neural warping procedure. The top, left panel illustrates the typical time sequence of sonar calls and echoes as a bat uses its sonar. The time from call onset to echo arrival varies as a function of object distance, and the time from call onset to the next call onset is actively adapted by the bat. This results in the window from echo arrival to call onset to be variable in time, precluding analyses that combine data across calls/conditions. To overcome this limitation, a time-warping procedure was employed to remap the proportional times of spikes from echo arrival to call onset into a standard time frame. The standard time frame was the average length of time for all echo-to-call windows. The right panel demonstrates how spikes times of variable length windows are warped to a standard time window. B. Real spike times for 18 different neurons (not all neurons spiked during this particular trial) for one echo-to-call window that is 200 milliseconds long. C. The average echo-to-call window length for this recording session was 62.5 milliseconds, and the real spikes times in D are therefore proportionally warped down to a window length of 62.5 milliseconds. D. Another echo-to-call window of the same 18 neurons activity, and the time duration of this window was 40 milliseconds. E. The real spikes times in D are therefore warped to a longer duration window of 62.5 milliseconds.

The time interval between echo arrival and call onset is variable due to the result of two phenomena: call interval is dynamically adapted depending upon sensory conditions and affects the time between echo arrival and call onset; and second, echo arrival times vary with the position of objects, and these positions change as the bat/object moves through space. This variability poses a challenge when averaging neural activity across multiple echo-to-call windows in a recording session – uniform window lengths are needed when averaging across multiple echo-to-call renditions. To overcome this issue in our data, we employed a time-warping approach to standardize the length of echo-to-call time windows within each recording session (see Fig 2). The warping procedure is described in detail in the methods, but in brief, for each recording session we first determine the average echo-to-call window length in time. We then calculate the proportional spike times in all of the original echo-to-call data set and warp the proportional times onto the average-length echo-to-call window. For instance, if single echo-to-call window in the original data is 100 milliseconds long with spike events at 50 milliseconds and 75 milliseconds, these time intervals would be converted into 50% and 75% respectively. The proportional spike time values are then mapped onto the corresponding time-points in the warped, average-length echo-to-call window. Figure 2 describes this in a schematic for two scenarios, 2b and 2c show an example of warping a longer original echo-to-call time interval (200ms) to a shorter average-length window (62ms). Fig 2d and 2e demonstrate this procedure in the context of warping a shorter original time interval (40ms) to a longer average-length echo-to-call time window (62ms). The time warping technique allows us to combine across multiple echo-to-call time windows that are variable in duration and examine average changes in population activity patterns.

To determine whether our pooled SC neurons show different patterns of activity during different behavioral events, we employed a dimensionality reduction technique on our warped, echo-to-call aligned data. We first computed instantaneous spike rates and then performed a principal component analysis (PCA) on the z-scored firing rates for each recording session (Fig 3). The input data to our PCA are neural spike rates, and the PCA output (i.e., scores) therefore identify the largest fluctuations in spike rates in the pooled neurons. Larger changes across PCA dimensions therefore denote larger relative changes in the firing rates across the recorded neurons. Example PCA outputs for four different recording sites are shown in Figure 3. Figure 3a shows data from a recording session (platform-tracking paradigm) where 15 different neurons were recorded, and five of those neurons’ activity patterns are shown in this panel (identified with a circled number). The representative neurons in Figure 3a show a variety of response types. Neuron 3 shows a burst of activity immediately following echo arrival (i.e., sensory), neuron 5 shows spiking activity following echo arrival as well as during production of a call (i.e., sensorimotor), while neuron 1 has sustained firing throughout the echo-to-call window. The inset 3D graphs in Figure 3 show the first 3 principal components of the data set. Figures 3a, 3b, 3c, and 3d show the average trajectory through PCA space for 4 different recording sessions. The green circle denotes the beginning of the neural vector through PCA space (i.e., when the echo arrives at the ears), and the red circle marks the end point (i.e., the onset of the upcoming call). The start and end of the neural PCA vectors trend towards similar locations in state space, while vectors diverge away from those locations through the middle of the echo-to-call window. The change in trajectories through PCA-space represent differences in the relative activity patterns across the pool of recorded neurons, likely related to changes in sensory to motor processing stages.

**Figure 3.**
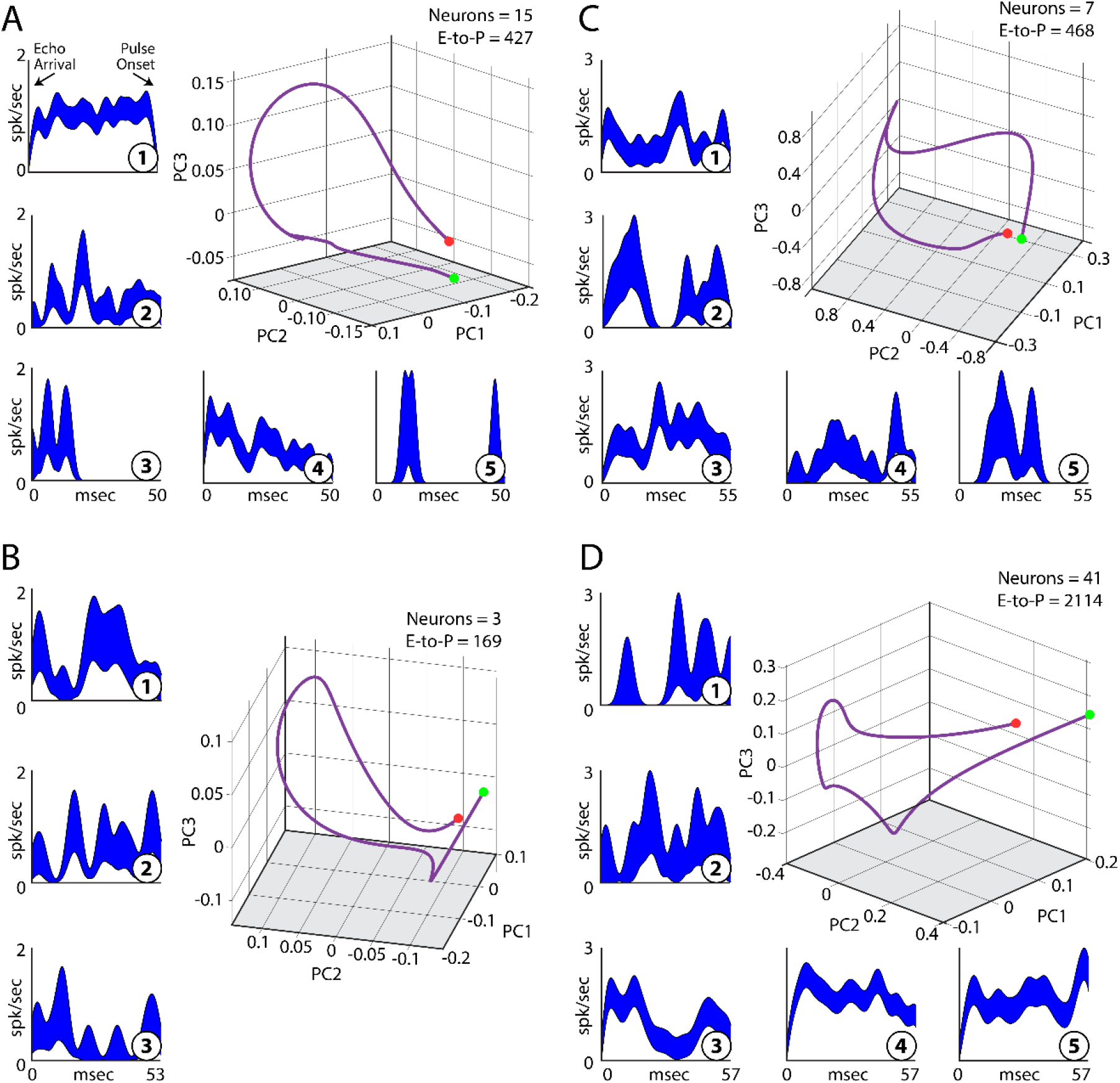
A.Left column and bottom row are PSTHs from 5 (numbers in circles) simultaneously recorded neurons during the echo-to-call window during the platform-tracking task. Plotted is the mean +/-s.e. of the spike rate. The 3D inset plot is the first 3 principal components from the dimensionality reduction performed on the entire data set which included 15 neurons, and 427 echo-to-call windows. The green circle marks the start of the trajectory (i.e., the time of echo arrival), while the red dot marks the end of the trajectory (i.e., the onset of the call). B. Another example as in A from a separate recording session with 3 neurons and 169 echo-to-call windows (platform-tracking task). C. Third example from a session with 7 neurons and 468 echo-to-call windows (free-flight task). D. Example from a recording session with 41 neurons and 2114 echo-to-call windows (free-flight task).

The neural trajectories reveal how simultaneously recorded neurons interact across behaviorally relevant windows of time. We can therefore analyze how pools of neurons change their coordinated firing rates under different conditions. We first asked if neuron-to-neuron interactions vary in the bat SC for calls produced at different rates. One of the main influences on call rate is target distance (Griffin, 1958), implying that changes in sensory signals about target range influence motor commands for call rate. We hypothesize that different patterns of SC sensory and motor activity occur for different rates of call production, resulting in different trajectories through PCA space. To test this prediction, we sorted the platform-tracking task data into four different call-interval bins (i.e., the inverse of call rate, Figure 4a, 0-50 milliseconds, 51-100 milliseconds, 101-150 milliseconds, and 151-200 milliseconds), and data from the free-flight task into 3 different call interval bins (Figure 4b, 1-30 milliseconds, 31-60 milliseconds, and 61-90 milliseconds). We then categorized the output scores of the PCA based upon these call interval bins. Two examples of this analysis are shown in Figures 4c and 4d, with Fig 4c showing example data from the platform-tracking task and 4d showing data from the free-flight task. The trend observed in these two examples (and confirmed in the larger data set, Figure 4e), is that longer call intervals result in longer trajectories through the neural state space. Since all echo-to-call windows have been warped to the same length regardless of the call interval, echo-to-call time duration does not explain the differences in trajectory length. For the platform-tracking task (Figure 4e, left), the trajectory is shortest for 0-50 millisecond call intervals (red), and then gets larger for each successive call interval bin (p < 0.05, 2-way repeated measures ANOVA). For the free-flight task data, we see a similar trend across the three call-interval bins: the trajectory length for the shortest call interval bin (1-30 msec) is significantly less than the larger bins (p < 0.05, 2-way repeated measures ANOVA). Importantly, the randomly selected *control* data (see Methods) had significantly shorter trajectories within each call-interval bin than the original data (p < 0.05, 2-way repeated measures ANOVA), and lacked a significant change in trajectory length as seen in the original data set. These results show that during the platform tracking task and free flight task, more rapid sensorimotor events are accompanied by shorter trajectories through the neural state space.

**Figure 4.**
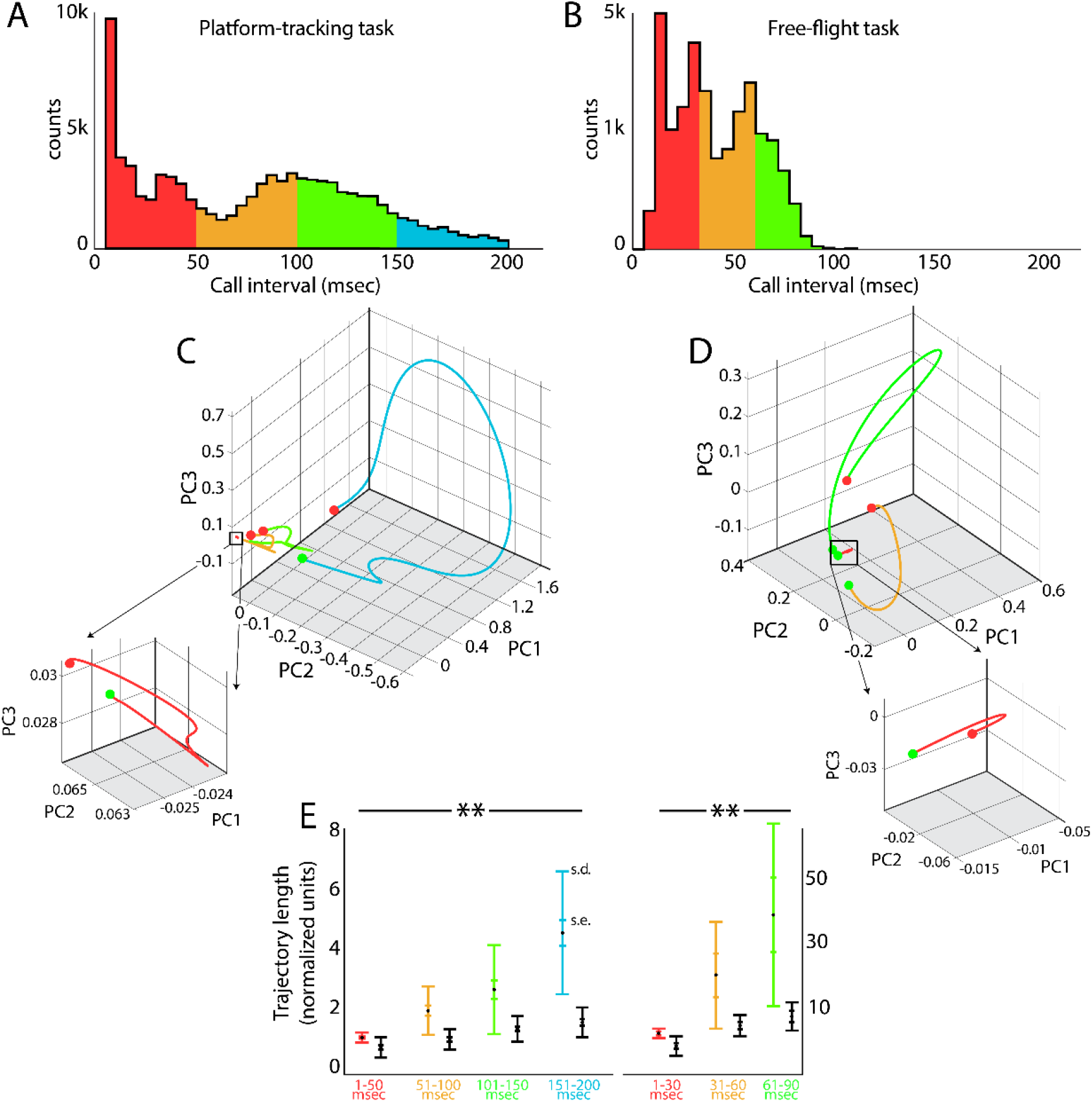
A.Histogram of the call interval distribution over all recordings, truncated at an interval of 200 milliseconds, for the platform tracking task. The call intervals are then split into 4 evenly spaced bins (color-coded). B. Histogram of the call interval distribution over all recordings, truncated at an interval of 200 milliseconds, for the free-flight task. The call intervals are then split into 3 evenly spaced bins (color-coded). C. One example from the platform-tracking task of the neural state space of SC activity sorted by the 4 call interval categories (same color scheme as in A) as outlined in panel A. Note the increase in trajectory length as call interval increases. D. One example from the free-flight task of the neural state space of SC activity sorted by the 4 call interval categories (same color scheme as in B) as outlined in panel A. Note the increase in trajectory length as call interval increases. E. Quantification of trajectory length for each call interval category for the platform-tracking task (left) and free-flight task (right). Randomly time-sampled windows are shown in black (see methods). A 2-way repeated measures ANOVA was used to test significance across conditions. There was a significant interaction between call interval bins (f = 124.81, p = 8.5e-10 for platform-tracking task; f = 146.67, p = 2.2e-10 for free-flight task), and a significant interaction between the original data set (colored bars) and the randomly selected data set (black bars) (f = 181.02, p = 3.6e-11 for platform-tracking task; f=201.13, p = 1.47e-11 for free-flight task).

In addition to trajectory length varying with call interval bins, the general shape of each trajectory also shows consistencies. The two examples shown in Fig 4c and 4d demonstrate how the trajectories start and end at nearby locations in state space, but deviate away from these locations during the middle portion of the trajectory. Prior work has shown that auditory response latencies in the SC are 5-30 milliseconds after the echo arrives at the ear (Valentine and Moss, 1997; Wohlgemuth and Moss, 2016), implying that the trajectories would start to deviate at a delay from echo arrival (the start of each trajectory, green circle). This feature of the data was quantified in Figure 5. For most call-interval bins (in platform tracking and free flight tasks), the distance in state space was shorter between the start and end of the trajectories than the distance from the start to the middle. This trend did not hold true for all comparisons (i.e., 51-100 msec bin for platform, 1-30 msec bin for free flight, p > 0.05 KS-test), but was found to be significant for the majority of call interval bins across both behavioral paradigms.

**Figure 5.**
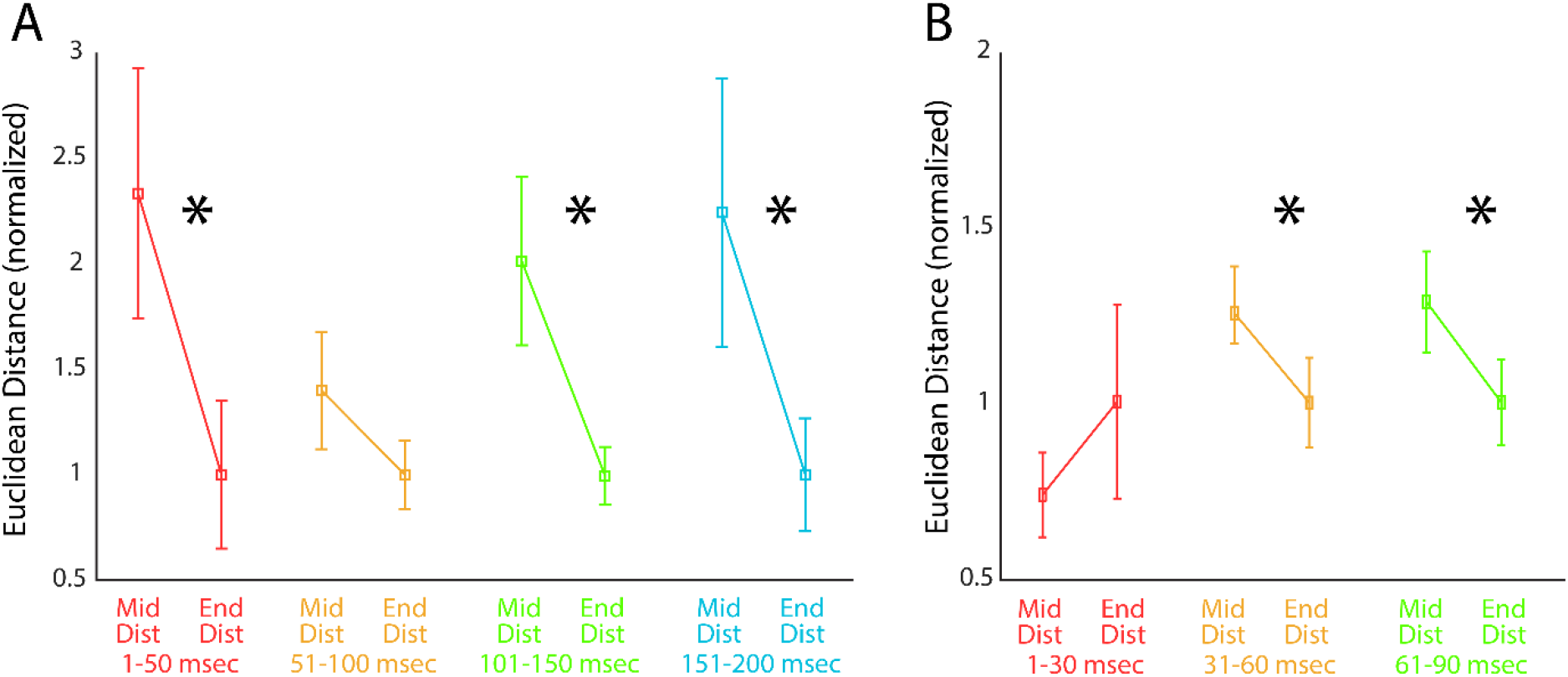
A.Data from the platform tracking task. Euclidian distances in n-dimensional space from the start of the trajectory to either the middle or end of the trajectory. Distances are normalized by the start-to-end distance. Data are color-coded as in Figure 5 with respect to call-interval bin. B. Data from the free-flight task across the three call-interval bins. Asterisks denote significance (p < 0.05, KS-test).

In addition to increases in call rate associated with decreasing object distance, bats also produce sonar sound groups (SSGs), or brief epochs of increased call rate, while they performed both behavioral tasks (see Fig 6a and 6b). Past work found SSGs are produced when bats fly in cluttered environments (Hulgard and Ratcliffe, 2016; Moss and Surlykke, 2010), which is similar to the free-flight task (Kothari et al, 2018). Other research shows an increase in SSG use when bats track erratically moving prey (Kothari et al., 2014), which is similar to aspects of the platform-tracking task. Analysis of echolocation audio data in these experiments revealed that about 54% of calls produced by bats in the free-flight task were SSGs (Fig 6c, right), and 20% of the calls produced by bats in the platform-tracking experiment were associated with SSGs (Fig 6c, left).

**Figure 6.**
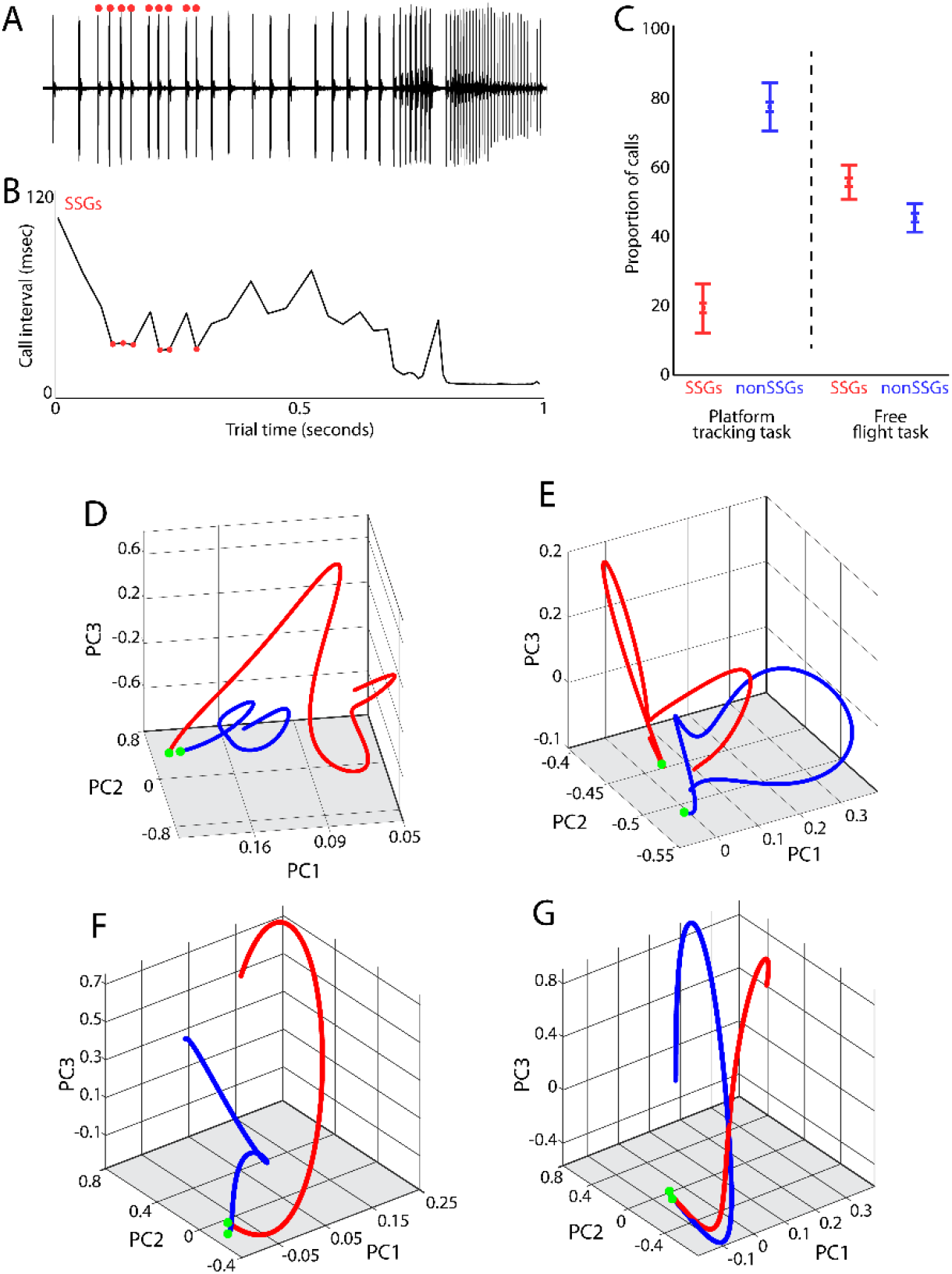
A.Example oscillogram from one trial of a bat tracking a moving target. Note both the decrease in call interval over time, as well as the brief decreases in call interval indicated by the red circles. B. The call intervals for the recording in panel A. The red circles indicate when the bat was producing sonar sound groups (SSGs). SSGs are defined as a small burst of calls at a shorter call interval. This is indicated as a quick drop in call interval surrounded by larger call intervals in panel B. C. The proportion of sonar calls that are found in SSGs versus those not in and SSG in the platform-tracking task (left) and free-flight task (right). D. Example neural state space trajectories for echo-to-call windows during SSG (blue, n=391) or non-SSG (red, n=111) production for one recording session during the platform-tracking task. The green circles mark the start of each trajectory. E. Another example of echo-to-call windows during SSG (blue, n=83) and non-SSG (red, n=57) for a separate recording session in the platform-tracking task. F. An example of the neural state space trajectories for data from the free-flight task showing SSG (blue, n=221) and non-SSG (red, n=92) state spaces for one recording session. G. Another neural state space example for SSG (blue, n=795) and non-SSG (red, n=312) echo-to-call windows during the free-flight task.

We analyzed coordinated SC activity in sensory-to-motor windows (i.e., echo-to-call) for SSG and non-SSG call sequences. Four examples of the differences in neural state between SSG production and non-SSG production are shown in Fig 6: 2 from the platform-tracking task (Fig 6d and 6e), and 2 from the free-flight task (Fig 6f and 6g). For each example, the SSG data is plotted in blue, and the non-SSG data is plotted in red. The start of each trajectory is marked with a green circle. Interestingly, the start locations of the SSG and non-SSG trajectories are in close proximity, but then begin to deviate later in the trajectory. As mentioned, SC auditory responses are delayed by as much as 30 milliseconds with respect to echo arrival (Valentine and Moss, 1997; Wohlgemuth and Moss, 2016). In the examples shown in Figure 6D-E, we see that SSG (blue) and non-SSG (red) neural PCA vectors begin to deviate soon after echo arrival and terminate at different locations in the PCA space (Figure 6), suggesting that SC activity diverges during sensory processing for SSG and nonSSG echoes.

To quantify how SC activity deviates for SSG and nonSSG call production, we measured the point-by-point distances in neural state space between the SSG and non-SSG trajectories for all recording sessions in one platform-tracking bat (Figure 7a, each line is one recording session, n=24). The average length of time from echo arrival to call onset in this bat was ∼70 milliseconds, and all neural windows are therefore warped to this time. The average of this data, as a percentage of time from echo arrival to call onset, is shown in Figure 7b in blue. The blue line plots the mean +/-s.e.m. for the data from the bat in Figure 7a; the purple line represents the data from the platform target tracking task, and the green line is the data from the free-flight task. The single bat data, as well as the combined bat data, quantitatively demonstrate the trend shown in Figure 6: neural states are similar at the time of echo arrival for both SSG and non-SSG data (0-10% bin), subsequently diverge (until the 50% mark from echo arrival to call onset), and then terminate at different locations in state space (90-100% bin). These trends are consistent across both behavioral paradigms.

**Figure 7.**
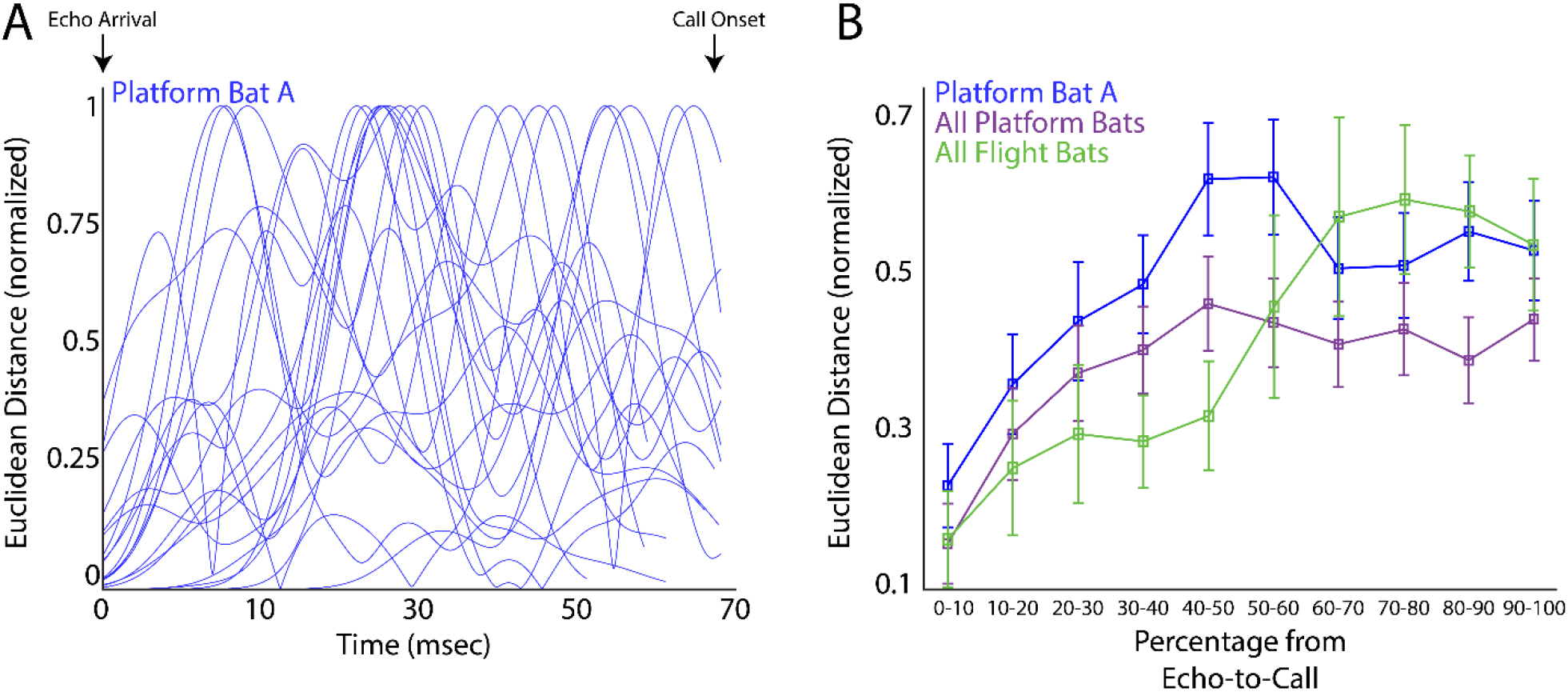
A.Normalized (i.e., 0 to 1) Euclidean distances in n-dimensional space between SSG and non-SSG trajectories through neural state space over echo-to-call windows. This is data across all recording sessions for one bat from the platform tracking task, and the average echo-to-call window time was 68 milliseconds for this bat. B. Binned average trajectory distances between SSG and non-SSG echo-to-call windows. The binning was done proportionally in 10% bins of total echo-to-call time for each bat so that data could be combined across bats. In blue is the data from the bat in panel A, in purple is that combine data across all bats in the platform-tracking task, and in green is combined data from all bats in the free-flight task. Plotted is the mean +/-s.e. normalized trajectory distance for each percentile bin.

SSGs are often produced in sequence, with multiple SSGs vocalized in a row. For instance, 2 call-SSGs (i.e., doublets) were often produced 3 times in sequence (6 calls in total) in both the platform tracking task and the free flight task. An example of the 3-SSG call pattern is shown in Figure 8a where the bat concatenated several SSGs in sequence. We analyzed changes in the neural PCA vectors for the first, second, and third SSG doublet in a sequence to determine whether the SC uses a different coding strategy for each SSG in a 3-SSG sequence. Examples of this analysis are shown in Figure 8b. For this analysis, neural PCA vectors are more complex than those in prior figures because the SSG windows include a pair of echo-to-call windows and three transitions (i.e., echo-call-echo-call), whereas prior figures show a single echo-to-call window and a single transition (i.e., echo-call). Figure 8b shows 3 examples of the neural state space trajectories for the first (blue), second (red), and third (green) doublet SSG in a 3-SSG sequence, 2 from bats trained on the platform-tracking task (top and middle), and one bat from the free-flight task (bottom). These examples demonstrate the overall trend: the length of neural state space vectors for successive SSGs decrease in length, with a significant difference between the 1^st^ and 3^rd^ SSG (Figure 8c, p < 0.001, permutation test). A control analysis was performed on series of 6 vocalizations in sequence that are nonSSG vocalizations (Figure 8a and 8c, purple lines) to determine if SSG production is different than nonSSG production in the SC. For this analysis, we analyzed neural PCA vector length across the first, second, and third pair of nonSSG vocalizations in a row (e.g., 6 calls in sequence that are not SSGs), and we did not find a decrease in trajectory length across the nonSSG vocal pairs. These data suggest that SSG production is categorically different than nonSSG production in terms of SC population activity, and this was true for data collected in the platform tracking and free flight tasks.

**Figure 8.**
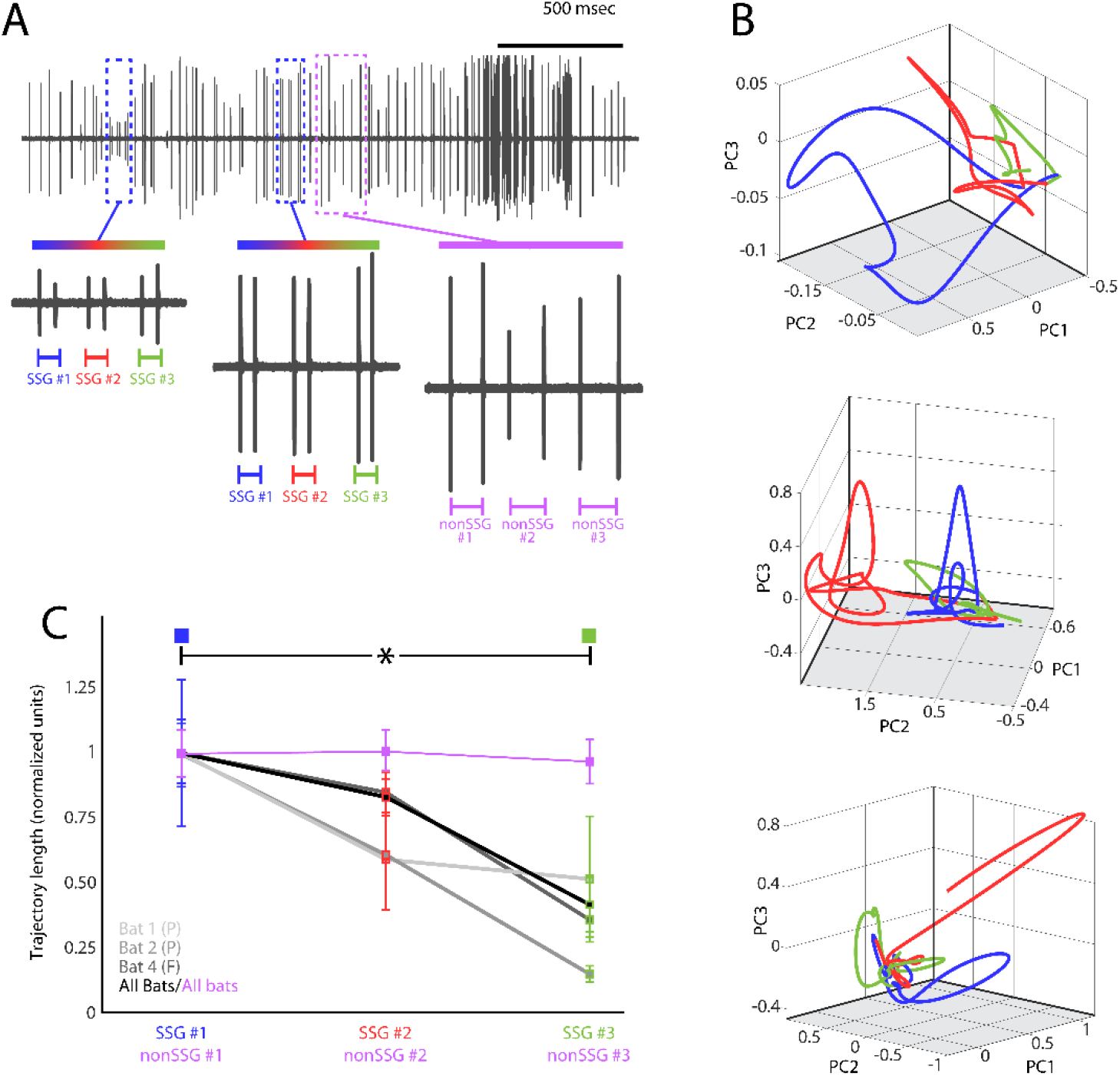
A.Example oscillogram of one vocal sequence from a bat tracking a prey item on a platform. Highlighted in red-green-blue colors are two instances of the bat producing 3, doublet SSGs in a row (blue = first, red = second, green = third) for each SSG in the sequence. In purple is a series of 6 vocalizations in sequence that are **not** SSGs that serves as a control sequence. **B.** Example neural state space trajectories for echo-to-call windows during SSG production. Single SSGs in blue (i.e., one pair of calls), double SSGs in red (i.e., two pairs of calls), triple SSGs in green (i.e., three pairs of calls). Top, Bat 1 from platform tracking experiment; middle, Bat 2 from platform tracking experiment; bottom, Bat 4 from free-flight task. C. Normalized, average trajectory lengths for single SSGs (blue), double SSGs (red), and triple SSGs (green). Data from individual bats are shown in greyscale, summary data across all bats in black. There is a significant decrease in trajectory length from single SSGs to triple SSGs (p < 0.001, permutation test). The purple line plots the change in trajectory length across the control vocal series in A of 6 nonSSG vocalizations in sequence.

## Discussion

We characterized changes in sensorimotor processing in the midbrain superior colliculus (SC) of echolocating bats engaged in sonar-guided target tracking and navigation tasks. We used a population-based approach to analyze multi-neuron recordings in conjuction with measurements of the bat’s sonar call rate and temporal patterning. By applying dimensionality reduction and analyzing sensorimotor dynamics in the resulting ‘neural sub-spaces’ of simultaneously recorded SC neurons, we identified several aspects of SC population activity that demonstrate its general role in adaptive active sensing behaviors. First, we found that as the bat increased its call rate, the trajectory length through neural subspace decreased. Second, we observed that pooled SC activity occupied distinct subspaces during echo arrival, sensory processing, and call production. Lastly, we found two interesting effects on pooled SC activity when the bat alters the temporal structure of its calls to produce sonar sound groups (SSGs) for increased spatial attention. We found that neural trajectories for SSG and nonSSG calls begin at similar state spaces, but diverge before call production. And we also found that when SSGs are produced repeatedly in sequence, the neural trajectories get progressively shorter. Together, these findings reveal shared neural dynamics in SC population activity across both spatial navigation and target-tracking behaviors. The importance of each finding is discussed below in more detail.

### Length of neural subspace trajectories indicate degree of sensorimotor integration

The length of a trajectory through neural state space reflects the magnitude of change in population activity. Because we applied principal component analysis (PCA) to the instantaneous firing rates of simultaneously recorded neurons, the resulting state space captures the largest relative changes in firing rates across the pooled neurons. A longer trajectory indicates greater variation in relative firing rates, suggesting that the neural population underwent more substantial dynamic changes during that period. In contrast, a shorter trajectory implies more stable or less variable firing patterns across the population. Thus, trajectory length provides a quantitative readout of how much neural population activity shifts over time in response to sensory inputs and in preparation for motor outputs.

We predicted that during very short call intervals, neural trajectories through state space would be constrained – resulting in shorter trajectories compared to those during longer call intervals. Our data support this prediction: as the call rate increases (call interval decreases), we observed a decrease in the length of neural trajectories. In contrast, control analysis using randomly selected (i.e. non-echo-to-call) time windows of matched durations do not show this trend. The control windows for analysis are not aligned to echo arrival and call onset, showing that the changes in trajectory length in the neural state space are indicative of active call rate based adjustments of sensorimotor processing and not an artifact of shortening the analysis window. Importantly, because some of the shortest call intervals span only a few milliseconds, our findings also reveal a potential processing constraint in terms of SC population dynamics – SC activity cannot change significantly when sensorimotor windows are very short. One solution is to integrate over longer windows of time when producing calls in rapid succession. Prior behavioral work supports this idea: bats tracking moving targets from a stationary position have been shown to integrate information over several echo-call pairs to construct internal models of target trajectories, which they use for adaptive motor control (Salles et al.,2020). Moreover, the shorter trajectories for higher call rates suggest that the SC quickly transitions from sensory to motor states at these instances. Past work has shown dynamic shifts in sensory and motor SC latencies when calls are produced at higher rates (Wohlgemuth et al, 2018), and the current work suggests that this may be mediated by activating a select pool of SC neurons with tuning profiles specific to the task.

### Sonar-guided spatial attention drives divergent trajectories through neural subspace

In addition to measuring overall trajectory length, we examined where in the trajectories any deviations occur. As shown in Figures 3-5, the trajectories begin and end at similar locations in neural state space, with a significant deviation emerging during the middle phase of the echo-to-call window. The close proximity of the start (sensory phase – echo arrival) and end (motor phase - call onset) points suggest that these events share a common underlying neural processing state. We propose that the SC resets its neural state at the end of each motor event (sonar call) to prepare for the next sensory event (echo arrival). This type of reset mechanism is consistent with prior modeling studies in human motor control, which have described similar dynamics during sequences of actions (Egger et al., 2020). We propose that deviations in the middle of the trajectories represent fluctuations in population-level activity indicative of the transition from sensory to motor processing. We find shorter trajectories for shorter call intervals, suggesting that the change in SC population activity from sensory processing to motor preparation is comparatively smaller. For longer call intervals, longer trajectories are observed indicating a larger change in the time window from sensory reception to motor production. Taken together, these results suggest that sensorimotor integration for longer call intervals involves larger changes in SC activity than calls produced in rapid succession. This finding relates back to the bat’s increase in call interval at larger target distances versus shorter call interval at reduced target distances. As described above, bats use echo information to dynamically adjust their call interval as they close in on a target (during the target tracking phase), but once they are in the immediate vicinity of the target (capture phase), call interval becomes very stereotyped (Moss and Surlykke, 2010). We therefore propose that the magnitude of deviation in neural state space (i.e. amount of change in pooled SC activity) over the echo-to-call window, is correlated with the degree of behavioral adaptation performed. Importantly, since we observe similar neural state space dynamics during in-flight navigation and target tracking behaviors, these population dynamics reflect a shared neural correlate of sensorimotor encoding across behaviors.

We also observed a divergence in neural trajectories when the bat adapted temporal patterning to produce SSGs (a burst of calls at a higher rate). Bats produce SSGs when flying in complex environments (Petrites et al 2009, Moss et al, 2006, Hulgard and Ratcliffe, 2016), when tracking erratically moving targets (Moss and Surlykke, 2010, Kothari et al, 2014, Kothari et al, 2018b), and when navigating around obstacles (Petrites et al, 2009, Kothari et al, 2018a). Thus, SSG calls are thought to be categorically different than nonSSG calls, such that SSGs might be an indirect measure of sonar guided attention (Moss and Surlykke, 2010, Kothari et al., 2018). To examine whether SC activity is different for SSG and nonSSG production, we compared trajectories through neural state space for each motor adaptation. We find that neural trajectories for upcoming SSG and nonSSG calls start at similar locations, and then begin to diverge partway through the echo-to-call window (Figure 7). This finding implies that the SC is in a relatively similar state before echo arrival for both call types, and then the arriving echo information either drives nonSSG call production or a behavioral shift to SSG production. These data suggest that SSG production involves a different SC activation pattern than nonSSG production, which is supported but earlier studies (see Kothari et al., 2018). Prior work reported that SC sensory neurons show significant shifts in their 3D spatial tuning profiles in reponse to echoes returning from SSGs compared to echoes returning from nonSSGs. Further, there was a concomitant increase in the power of the gamma band of the LFP during SSG production. Our data also show significant deviations in pooled SC activity for SSG production, lending further evidence to the categorically different motor control program for this call pattern. Interestingly, our data show that for orienting behaviors towards the same location in space, different sensing strategies (SSG and nonSSG calls) recruit different SC activation patterns. This finding is interesting in that the SC is mostly thought to have a space code, but we find that different behaviors executed to the same spatial location can differentially activate the SC. Importantly, we find this for both the free-flight and the platform-tracking tasks.

### Adaptive temporal patterning suggests sensorimotor refinement over time

When bats fly in cluttered environments, they attend to objects along their flight path, and the SC representation of object depth is sharpened as bats produce SSG calls to closely inspect the object (Kothari et al, 2018). Based on these prior findings, we hypothesized that repeated SSG production serves to reduce sensory uncertainty, which would be reflected as a shortening of neural trajectory length over successive SSGs in sequence. We analyzed the trajectories through neural state space for 3-pairs of consecutive SSGs (i.e., SSG #1, SSG #2, and SSG #3). The data support our hypothesis and show a decrease in the length of neural subspace trajectories for each successive SSG, and a significant difference between SSG #1 and SSG #3. Importantly, control sequences of 3-pairs of consecutive nonSSG calls in a sequence showed no changes in subspace trajectories. We propose that as sensory evidence is accumulated and integrated across the sequence of SSGs, this results in a reduction in sensorimotor processing across successive SSGs. In our data, this is represented as a progressive decrease in trajectory length in neural state space, with reduced trajectory length serving as a readout of the refinement of sensory information and reduced sensory uncertainty in the brain. These results support the finding of sharpening of depth tuning of SC neurons with SSG production (Kothari et al., 2018).

## Conclusions

In this study, we applied PCA-based dimensionality reduction techniques to analyze sensorimotor population dynamics in the bat superior colliculus (SC). Our goal was to determine how variations in sensorimotor processing are linked to distinct behavioral adaptations. We identified significant differences in SC population activity associated with different vocal behaviors, along with evidence of sensorimotor refinement signals. These findings suggest that the SC plays a key role in integrating sensory inputs with motor planning. Future investigations should explore how distinct functional states within sensory neuron populations influence motor preparation.

## BIBLIOGRPAHY

Bernardin, D., Kadone, H., Bennequin, D., Sugar, T., Zaoui, M., and Berthoz, A. (2012). Gaze anticipation during human locomotion. Exp. Brain Res. 223, 65–78. doi:10.1007/s00221-012-3241-2.

Churchland, M. M., Cunningham, J. P., Kaufman, M. T., Foster, J. D., Nuyujukian, P., Ryu, S. I., et al. (2012). Neural population dynamics during reaching. Nature 487, 51–56. doi:10.1038/nature11129.

Churchland, M. M., Yu, B. M., Ryu, S. I., Santhanam, G., and Shenoy, K. V (2006). Neural Variability in Premotor Cortex Provides a Signature of Motor Preparation. J. Neurosci. 26, 3697–3712. doi:10.1523/JNEUROSCI.3762-05.2006.

Churchland, M. M., Yu, B. M., Sahani, M., and Shenoy, K. V (2007). Techniques for extracting single-trial activity patterns from large-scale neural recordings. Curr. Opin. Neurobiol. 17, 609–618. doi:10.1016/j.conb.2007.11.001.

Cynader, M., and Berman, N. (1972). Receptive-Field Organization of Monkey Superior Colliculus. J. Neurophysiology 35, 187–201. Available at: http://jn.physiology.org/content/jn/35/2/187.full.pdf [Accessed May 1, 2017].

Egger, S. W., Le, N. M., and Jazayeri, M. (2020). A neural circuit model for human sensorimotor timing. Nat. Commun. 11, 3933. doi:10.1038/s41467-020-16999-8.

Falk, B., Jakobsen, L., Surlykke, A., and Moss, C. F. (2014). Bats coordinate sonar and flight behavior as they forage in open and cluttered environments. J. Exp. Biol. 217, 4356–64. doi:10.1242/jeb.114132.

Gandhi, N. J., and Katnani, H. A. (2011). Motor functions of the superior colliculus. Annu. Rev. Neurosci. 34, 205–31. doi:10.1146/annurev-neuro-061010-113728.

Ghose, K., and Moss, C. F. (2003). The sonar beam pattern of a flying bat as it tracks tethered insects. J. Acoust. Soc. Am. 114, 1120–31. Available at: http://www.pubmedcentral.nih.gov/articlerender.fcgi?artid=3384009&tool=pmcentrez&rendertype=abstract [Accessed September 10, 2015].

Goldberg, M. E., and Wurtz, R. H. (1972). Activity of superior colliculus in behaving monkey. I. Visual Receptive Fields of Single Neurons. J. Neurophysiol. 35, 542–559.

Griffin, D. (1958). Listening in the dark: the acoustic orientation of bats and men. Yale University Press.

Griffin, D. R., Webster, F. A., and Michael, C. R. (1960). The echolocation of flying insects by bats. Anim. Behav. 8, 141–154. doi:10.1016/0003-3472(60)90022-1.

Hulgard, K., and Ratcliffe, J. M. (2016). Sonar sound groups and increased terminal buzz duration reflect task complexity in hunting bats. Sci. Rep. 6, 21500. doi:10.1038/srep21500.

Jakobsen, L., Olsen, M. N., and Surlykke, A. (2015). Dynamics of the echolocation beam during prey pursuit in aerial hawking bats. Proc. Natl. Acad. Sci. U. S. A. 112, 8118–23. doi:10.1073/pnas.1419943112.

Jakobsen, L., Ratcliffe, J. M., and Surlykke, A. (2013). Convergent acoustic field of view in echolocating bats. Nature 493, 93–6. doi:10.1038/nature11664.

Kothari, N. B., Wohlgemuth, M. J., Hulgard, K., Surlykke, A., and Moss, C. F. (2014). Timing matters: sonar call groups facilitate target localization in bats. Front. Physiol. 5, 168. doi:10.3389/fphys.2014.00168.

Kothari, N. B., Wohlgemuth, M. J., and Moss, C. F. (2018). Dynamic representation of 3D auditory space in the midbrain of the free-flying echolocating bat. Elife 7, e29053. doi:10.7554/eLife.29053.

Leonardo, A. (2004). Experimental test of the birdsong error-correction model. Proc. Natl. Acad. Sci. 101, 16935–16940. doi:10.1073/pnas.0407870101.

Luo, J., Kothari, N. B., and Moss, C. F. (2017). Sensorimotor integration on a rapid time scale. Proc. Natl. Acad. Sci. 114, 6605–6610. doi:10.1073/PNAS.1702671114.

Ma, T. P., Cheng, H.-W., Czech, J. A., and Rafols, J. A. (1990). Intermediate and deep layers of the macaque superior colliculus: A golgi study. J. Comp. Neurol. 295, 92–110. doi:10.1002/cne.902950109.

May, P. J. (2006). “The mammalian superior colliculus: laminar structure and connections,” in, 321–378. doi:10.1016/S0079-6123(05)51011-2.

McIlwain, J. T. (1982). Lateral spread of neural excitation during microstimulation in intermediate gray layer of cat’s superior colliculus. J. Neurophysiol. 47. Available at: http://jn.physiology.org/content/47/2/167.long [Accessed April 9, 2017].

McIlwain, J. T., Albus, K., Appell, P. P., Behan, M., Becker, W., Jürgens, R., et al. (1991). Distributed spatial coding in the superior colliculus: A review. Vis. Neurosci. 6, 3–13. doi:10.1017/S0952523800000857.

Moss, C. F., and Surlykke, A. (2001). Auditory scene analysis by echolocation in bats. J. Acoust. Soc. Am. 110, 2207. doi:10.1121/1.1398051.

Moss, C. F., and Surlykke, A. (2010). Probing the natural scene by echolocation in bats. Front. Behav. Neurosci. 4. doi:10.3389/fnbeh.2010.00033.

Moss, C. F., Chiu, C., and Surlykke, A. (2011). Adaptive vocal behavior drives perception by echolocation in bats. Curr. Opin. Neurobiol. 21, 645–52. doi:10.1016/j.conb.2011.05.028.

Nelson, M. E., and MacIver, M. A. (2006). Sensory acquisition in active sensing systems. J. Comp. Physiol. A. Neuroethol. Sens. Neural. Behav. Physiol. 192, 573–86. doi:10.1007/s00359-006-0099-4.

Peck, C. K. (1990). Neuronal activity related to head and eye movements in cat superior colliculus. J. Physiol. 421, 79–104. doi:10.1113/jphysiol.1990.sp017934.

Petrites, A. E., Eng, O. S., Mowlds, D. S., Simmons, J. A., and DeLong, C. M. (2009). Interpulse interval modulation by echolocating big brown bats (Eptesicus fuscus) in different densities of obstacle clutter. J. Comp. Physiol. A. Neuroethol. Sens. Neural. Behav. Physiol. 195, 603–17. doi:10.1007/s00359-009-0435-6.

Quiroga, R. Q., Nadasdy, Z., and Ben-Shaul, Y. (2004). Unsupervised Spike Detection and Sorting with Wavelets and Superparamagnetic Clustering. Neural Comput. 16, 1661–1687. doi:10.1162/089976604774201631.

Roucoux, A., Guitton, D., and Crommelinck, M. (1980). Stimulation of the superior colliculus in the alert cat. II. Eye and head movements evoked when the head is unrestrained. Exp. brain Res. 39, 75–85. Available at: http://www.ncbi.nlm.nih.gov/pubmed/7379887 [Accessed February 16, 2017].

Russo, A. A., Khajeh, R., Bittner, S. R., Perkins, S. M., Cunningham, J. P., Abbott, L. F., et al. (2020). Neural Trajectories in the Supplementary Motor Area and Motor Cortex Exhibit Distinct Geometries, Compatible with Different Classes of Computation. Neuron 107, 745-758.e6. doi:10.1016/j.neuron.2020.05.020.

Sändig, S., Schnitzler, H.-U., and Denzinger, A. (2014). Echolocation behaviour of the big brown bat (Eptesicus fuscus) in an obstacle avoidance task of increasing difficulty. J. Exp. Biol. 217, 2876–84. doi:10.1242/jeb.099614.

Schroeder, C. E., Wilson, D. A., Radman, T., Scharfman, H., and Lakatos, P. (2010). Dynamics of Active Sensing and perceptual selection. Curr. Opin. Neurobiol. 20, 172–6. doi:10.1016/j.conb.2010.02.010.

Shenoy, K. V, Sahani, M., and Churchland, M. M. (2013). Cortical Control of Arm Movements: A Dynamical Systems Perspective. Annu. Rev. Neurosci. 36, 337–359. doi:10.1146/annurev-neuro-062111-150509.

Sober, S. J., Wohlgemuth, M. J., and Brainard, M. S. (2008). Central contributions to acoustic variation in birdsong. J. Neurosci. 28, 10370–9. doi:10.1523/JNEUROSCI.2448-08.2008.

Sparks, D. L. (1986). Translation of sensory signals into commands for control of saccadic eye movements: role of primate superior colliculus. Physiol. Rev. 66, 118–171.

Surlykke, A., Boel Pedersen, S., and Jakobsen, L. (2009). Echolocating bats emit a highly directional sonar sound beam in the field. Proc. Biol. Sci. 276, 853–60. doi:10.1098/rspb.2008.1505.

Surlykke, A., and Moss, C. F. (2000). Echolocation behavior of big brown bats, Eptesicus fuscus, in the field and the laboratory. J. Acoust. Soc. Am. 108, 2419–29. Available at: http://www.ncbi.nlm.nih.gov/pubmed/11108382 [Accessed September 10, 2015].

Valentine, D. E., and Moss, C. F. (1997). Spatially selective auditory responses in the superior colliculus of the echolocating bat. J. Neurosci. 17, 1720–33. Available at: http://www.ncbi.nlm.nih.gov/pubmed/9030631 [Accessed September 10, 2015].

Valentine, D. E., Sinha, S. R., and Moss, C. F. (2002). Orienting responses and vocalizations produced by microstimulation in the superior colliculus of the echolocating bat, Eptesicus fuscus. J. Comp. Physiol. A. Neuroethol. Sens. Neural. Behav. Physiol. 188, 89–108. doi:10.1007/s00359-001-0275-5.

Wohlgemuth, M. J., Kothari, N. B., and Moss, C. F. (2018). Functional organization and dynamic activity in the superior colliculus of the echolocating bat, Eptesicus fuscus. J. Neurosci. 38, 245–256. doi:10.1523/JNEUROSCI.1775-17.2017.

Wohlgemuth, M. J., and Moss, C. F. (2016). Midbrain auditory selectivity to natural sounds. Proc. Natl. Acad. Sci. U. S. A. 113. doi:10.1073/pnas.1517451113.

Wohlgemuth, M., Kothari, N., and Moss, C. (2017). Midbrain functional organization and dynamic coding for natural orientation. J. Neurosci. in revisio.

Wurtz, R. H., and Albano, J. E. (1980). VISUAL-MOTOR FUNCTION OF THE PRIMATE SUPERIOR COLLICULUS1. Ann. Rev. Neurosci 3, 189–226. Available at: http://www.annualreviews.org/doi/pdf/10.1146/annurev.ne.03.030180.001201 [Accessed March 10, 2017].

Wurtz, R. H., and Goldberg, M. E. (1971). Superior colliculus cell responses related to eye movements in awake monkeys. Science 171, 82–4. Available at: http://www.ncbi.nlm.nih.gov/pubmed/4992313 [Accessed February 16, 2017].

Zimnik, A. J., and Churchland, M. M. (2021). Independent generation of sequence elements by motor cortex. Nat. Neurosci. 24, 412–424. doi:10.1038/s41593-021-00798-5.

